# The dynamic proteome of influenza A virus infection identifies M segment splicing as a host range determinant

**DOI:** 10.1101/438176

**Authors:** Boris Bogdanow, Katrin Eichelbaum, Anne Sadewasser, Xi Wang, Immanuel Husic, Katharina Paki, Martha Hergeselle, Barbara Vetter, Jingyi Hou, Wei Chen, Lüder Wiebusch, Irmtraud M. Meyer, Thorsten Wolff, Matthias Selbach

**Affiliations:** Max Delbrück Center for Molecular Medicine, Robert-Rössle-Str. 10, 13125 Berlin; Unit 17 “Influenza and other Respiratory Viruses”, Robert Koch Institut, Seestr. 10, 13353 Berlin, Germany; Division of Theoretical Systems Biology, German Cancer Research Center, 69120 Heidelberg, Germany; Labor für Pädiatrische Molekularbiologie, Charité Universitätsmedizin Berlin, Augustenburger Platz 1, 13353 Berlin, Germany; Department of Biology, Southern University of Science and Technology, Xuanyuan Rd. 1088, 518055 Shenzhen, China; Charité Universitätsmedizin Berlin, 10117 Berlin, Germany

## Abstract

A century ago, influenza A virus (IAV) infection caused the 1918 flu pandemic and killed an estimated 20-40 million people. Pandemic IAV outbreaks occur when strains from animal reservoirs acquire the ability to infect and spread among humans. The molecular details of this species barrier are incompletely understood. We combined metabolic pulse labeling and quantitative shotgun proteomics to globally monitor protein synthesis upon infection of human cells with a human-and a bird-adapted IAV strain. While production of host proteins was remarkably similar, we observed striking differences in the kinetics of viral protein synthesis over the course of infection. Most importantly, the matrix protein M1 was inefficiently produced by the bird-adapted strain at later stages. We show that impaired production of M1 from bird-adapted strains is caused by increased splicing of the M segment RNA to alternative isoforms. Experiments with reporter constructs and recombinant influenza viruses revealed that strain-specific M segment splicing is controlled by the 3’ splice site and functionally important for permissive infection. Independent *in silico* evidence shows that avian-adapted M segments have evolved different conserved RNA structure features than human-adapted sequences. Thus, our data identifies M segment RNA splicing as a viral determinant of host range.

## INTRODUCTION

Influenza A viruses (IAVs) are negative-sense, single-stranded RNA viruses with a segmented genome. IAV infection causes seasonal epidemics and sporadically pandemic outbreaks in the human population with significant morbidity, mortality and economic burden. IAVs can infect both mammals (e.g. humans, pigs, horses) and birds (e.g. chicken, waterfowl). However, strains that are replicating in birds typically do not infect mammals and *vice versa*. Pandemics occur when influenza strains of avian origin with novel antigenicity acquire the ability to transmit among humans ^1^. Understanding the molecular basis of host specificity is therefore of high medical relevance.

The species barriers that hinder most avian IAVs from successfully infecting humans are effective at several steps in the viral life cycle. For example, the avian virus receptor hemagglutinin (HA) recognizes oligosaccharides containing terminal sialic acid (SA) that are linked to galactose by α2,3 ^2^. In the human upper respiratory airway epithelium the dominant linkage is of α2,6 type, to which human-adapted hemagglutinin binds. Despite these differences in receptor binding, many avian viruses are internalized by human cells and initiate expression of the viral genome. Such infections typically lead to an abortive, nonproductive outcome in human cell lines ^3–6^. Our understanding of this intracellular restriction is still incomplete. One well-established factor is the influenza RNA dependent RNA polymerase (RdRp): This enzyme catalyzes replication of the viral genome and transcription of viral mRNAs ^7^. Polymerases from avian strains are considerably less active in mammalian cells than their counterparts from mammalian-adapted strains ^8,9^. A wealth of experimental data described adaptive mutations that alter receptor specificity or fusion activity of HA (reviewed in ^10^) and polymerase activity (reviewed in ^11^). However, relatively little is known about the contribution of other Influenza A virus genes for permissive versus non-permissive infection ^12^.

A crucial aspect for permissive infection is the correct timing of viral gene expression: IAV proteins are produced at the specific phase of infection when they are needed ^13^. One example is the M gene, which encodes predominantly two polypeptides: The larger protein, M1, is produced from a collinear transcript. The smaller one, M2, is encoded by a differentially spliced transcript ^14^. M1 is the matrix protein with multiple functions that encapsulates the viral genome and also mediates nuclear export ^15,16^. M2 is a proton-selective channel that is an integral part of the viral envelope ^17,18^. The ratio of spliced to unspliced products increases during infection ^19^, which reflects the changing demands required for optimal viral replication.

Systems-level approaches have provided important insights into the molecular details of host-virus interaction ^20^. For example, RNAi screens identified host factors required for IAV replication ^21–23^. Also, interaction proteomics experiments identified many cellular binding partners of IAV proteins ^24–26^. A number of studies also quantified changes in protein abundance ^27–32^. However, these steady-state measurements cannot reveal the dynamic changes in protein synthesis during different phases of infection. Early studies used radioactive pulse labeling to monitor protein synthesis in IAV infected cells ^6,33^. However, radioactive pulse labelling cannot provide kinetic profiles for individual proteins. More recently, stable isotope labelling by amino acids in cell culture (SILAC) emerged as a powerful means to study the dynamic proteome ^34^. SILAC-based pulse labelling methods such as pulse SILAC (pSILAC) and dynamic SILAC can quantify protein synthesis and degradation on a proteome-wide scale ^35,36^. Moreover, metabolic incorporation of bioorthogonal amino acids such as azidohomoalanine (AHA) provides a means to biochemically enrich for newly synthesized proteins ^37^. In combination with SILAC, AHA labeling can be used to quantify proteome dynamics with high temporal resolution ^38–41^.

Here, we used metabolic pulse labeling and quantitative mass spectrometry to compare proteome dynamics upon infection of human cells with a human-adapted and a bird-adapted IAV strain. We found that host proteins behaved surprisingly similar but observed striking differences in the production of viral proteins, especially for the matrix protein M1. Follow-up experiments with reporter constructs, *in silico* studies and reverse genetics identified an evolutionarily conserved cis-regulatory element in the M segment as a novel host range determinant.

## RESULTS

### Quantifying the dynamic proteome of permissive and non-permissive IAV infection

To assess species specificity of influenza A viruses (IAVs) we used a model system comparing a low-pathogenic avian H3N2 IAV (A/Mallard/439/2004 - Mal) to a seasonal human IAV isolate of the same subtype (A/Panama/2007/1999 - Pan). While the avian virus is not adapted to efficient growth in cultured human cells and causes a non-permissive infection, the seasonal human virus replicates efficiently. We demonstrated previously that the Pan virus produces >1,000 fold more infectious viral progeny than the non-adapted virus, even though both strains efficiently enter human cells and initiate their gene expression program ^32^.

We reasoned that comparing the kinetics of protein synthesis upon infection with both strains might reveal determinants of species specificity. To this end we performed proteome-wide comparative pulse-labeling experiments by combining labeling with azidohomoalanine (AHA) and SILAC (Stable Isotope Labeling of Amino acids in Cell culture) (Figure 1A): Cells incorporate AHA instead of methionine into newly synthesized proteins when the cell culture medium is supplemented with this bioorthogonal amino acid ^42^. AHA contains an azido-group which can be used to covalently coupled AHA-containing proteins to alkyne beads via click-chemistry. In this manner, newly synthesized proteins can be selectively enriched from the total cellular proteome. Combining AHA labeling with SILAC reveals the kinetics of protein synthesis with high temporal resolution ^39,41^. First, we fully labeled human lung adenocarcinoma cells (A549) using SILAC. Second, individual cell populations were infected with either Pan or Mal virus or left uninfected. Third, all cells were pulse labeled with AHA for four hours during different time intervals post infection (0-4, 4-8, 8-12 and 12-16 hrs). The three cell populations for every time interval were then combined, lysed, and AHA-containing proteins were enriched from the mixed lysate using click chemistry (Figure 1B). After on-bead digestion, peptide samples were analyzed by high resolution shotgun proteomics.

**Figure 1.**
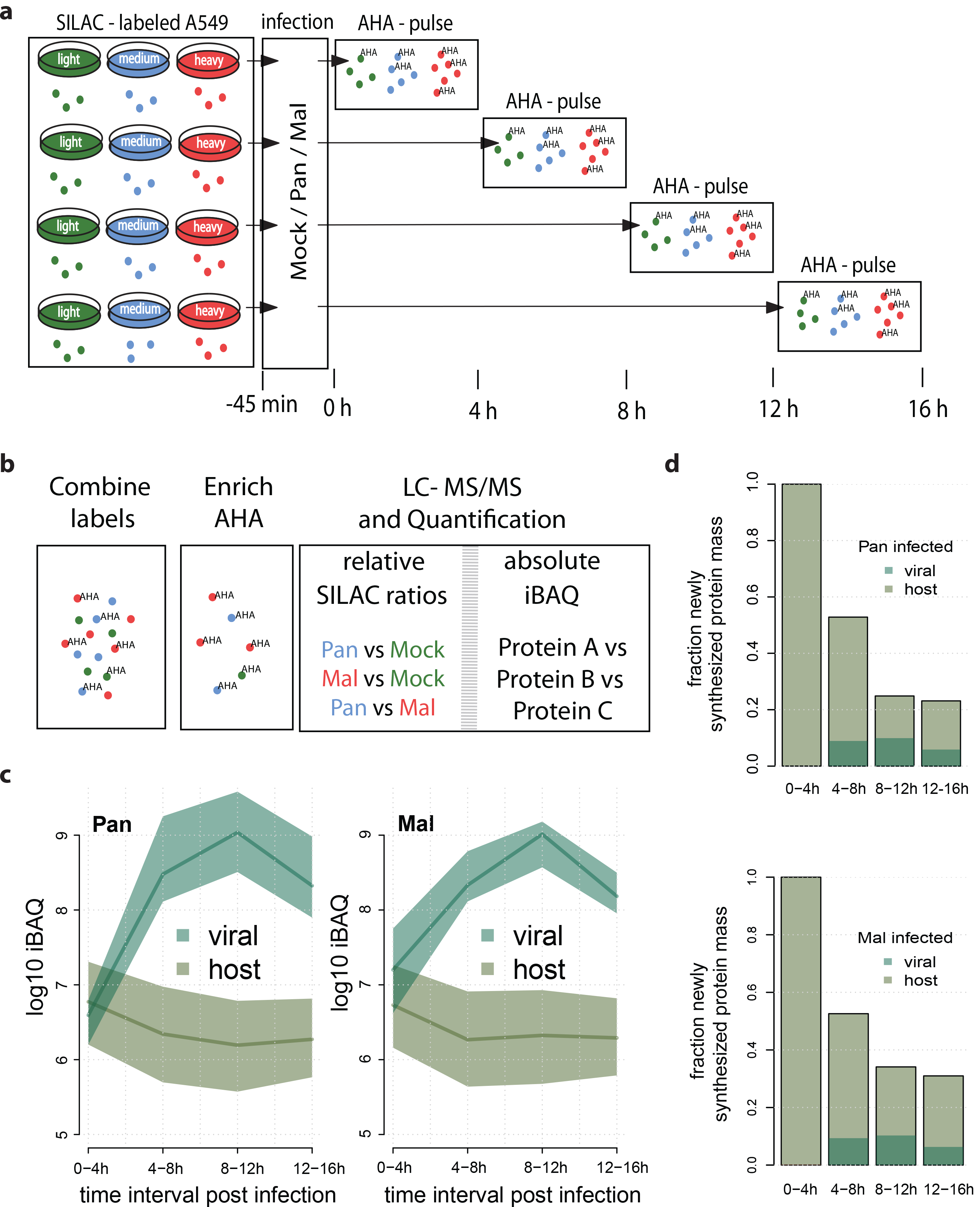
A strategy to quantify protein *de novo* synthesis proteome-wide. **a** SILAC L/M/H - labeled A549 cells were infected with the human seasonal H3N2 IAV isolate Panama (Pan) or the avian H3N2 isolate Mallard (Mal) or left uninfected. The methionine analogous AHA was given to the methionine-depleted medium in different 4 h intervals. **b** After lysis and enrichment for proteins that incorporated AHA, samples were subjected to shotgun-proteomics. Absolute and relative protein synthesis profiles were quantified for host and viral proteins. **c** iBAQ based quantification of protein synthesis levels for host and viral proteins in cell infected with either Pan or Mal virus, as indicated. Median, 25th and 75th percentile of the respective populations are given. **d** Quantification of the total newly synthesized protein mass for host and viral proteins with either Pan or Mal infection as indicated. Data was normalized to the 0-4 h time period.

We quantified proteins using two readouts: (i) SILAC-based relative quantification to assess differences in *de novo* protein synthesis and (ii) intensity-based absolute quantification (iBAQ) to quantify absolute amounts of newly synthesized proteins ^43^. Our data thus provides kinetic profiles for relative and absolute differences in *de novo* protein synthesis across the course of infection (Supplementary Table S1). In total, we identified 7,189 host and 10 viral proteins and quantified 6,019 proteins in at least two biological replicates with overall good reproducibility (Supplementary Figure 1).

### The dynamic host proteome

It is well established that IAV induces a global reduction in the production of host proteins. This host shutoff was attributed to a plethora of viral effector functions ^44^. To assess the host shutoff in our proteomic data, we investigated iBAQ values for viral and host proteins. As expected, viral proteins were potently induced while the production of host proteins decreased over time (Figure 1C-D). The difference between host and viral protein synthesis reached several orders of magnitude and was highest during the 8-12 h pulse interval. Moreover, the total cellular protein output dropped to ~24 % (Pan) or ~30 % (Mal) at later stages of infection, of which ~20-40 % was of viral origin (Figure 1E-F). At this level of detail, we observed no major differences between both strains. Thus, both strains initiate viral protein synthesis and induce the shutoff of host protein synthesis to an overall similar extent.

Next, we investigated the profiles of individual host proteins across the course of infection. For this, we directly looked at SILAC ratios comparing infected and non-infected cells (Figure 2A). As expected, synthesis of the vast majority of host proteins markedly decreased over time. However, some proteins were less affected by the host shutoff and displayed only a mildly decreased or even increased production. To assess this observation more systematically, we selected the proteins that were least affected by the shutoff at different pulse periods and performed gene ontology (GO) analysis. The heatmap of enriched GO terms provides a global overview of biological processes as the infection progresses (Figure 2B and Supplementary Table S2). For example, many well-known interferon-induced antiviral defense proteins (e.g. MX1, several IFIT proteins, several oligoadenylate synthase proteins) were relatively strongly produced at late stages of infection. Also, many ribosomal proteins (GO term “peptide chain elongation”) largely escaped the host shutoff. Interestingly, we also observed significant enrichment of proteins involved in steroid metabolism and mitochondrial proteins (mito-ribosomal, respiratory chain proteins) at early and intermediate stages of infection, respectively. Cellular responses to infection with the Pan and Mal strain were overall similar. To assess potential differences between permissive and non-permissive infection we compared protein log2 fold changes between both viruses directly (Figure 2C). Interestingly, type I interferon response proteins were first preferentially produced during non-permissive infection. At later stages, however, infection with the Pan virus elicited a stronger interferon response.

**Figure 2.**
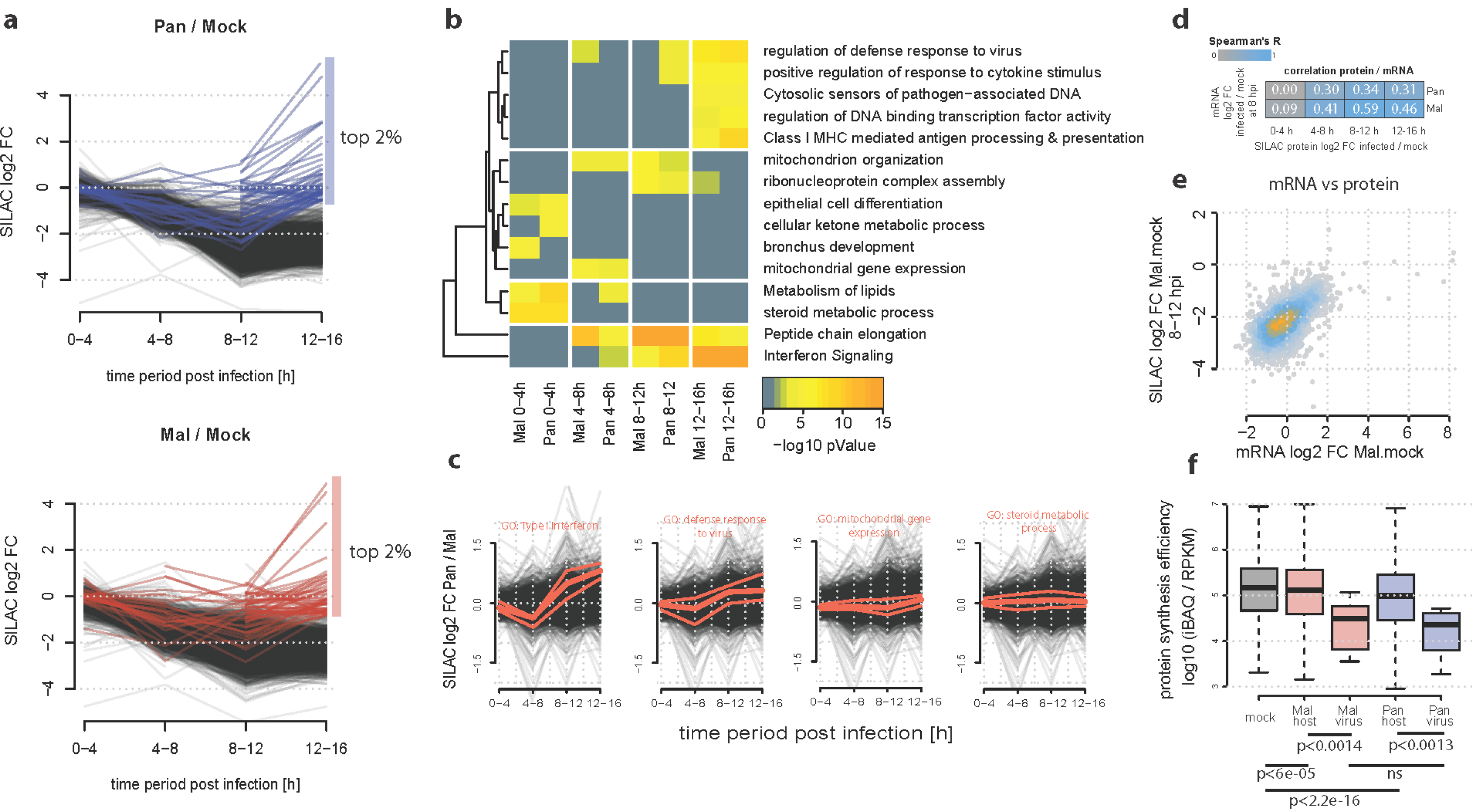
The dynamic host proteome of IAV infected cells. **a** SILAC profiles of host proteins infected with Pan (upper panel) or Mal (lower panel). The top 2% of proteins that are strongest produced during the last time period are highlighted. **b** Functional protein clusters least affected by host shutoff were identified. The two percent proteins with the highest ratios were selected for each time period and multi-list GO enrichment was performed. **c** Median, 25th and 75th percentile of proteins related to a selected set of GO terms are highlighted in the direct SILAC comparison Pan/ Mal. All other protein profiles in black. **d** Correlation coefficients when comparing the indicated RNA and protein level data. **e** Scatterplot of mRNA changes versus protein synthesis level changes. Blue and orange coloring of data points reflects the density of data points. **f** Protein synthesis efficiencies calculated from absolute mRNA (RPKM) and protein data (iBAQ). Data points outside the whiskers were removed for visibility. For the assessment of statistical significance wilcoxon rank sum tests were performed and p-values are given (ns: non-significant).

Several different hypotheses were made to explain the IAV induced host shutoff. This included mechanisms at the transcriptional ^45^, post-transcriptional ^46,47^ and translational ^48,49^ level. To study the relationship of mRNA and protein levels we quantified mRNA levels at 8 hours post infection by RNA-seq. mRNA level differences at this time point showed good correlation with corresponding differences in *de novo* protein synthesis, particularly during the subsequent 8-12 h period (Figure 2D-E). Thus, mRNA level changes play an important role for the shutoff of individual mRNAs, corroborating the view that the host shutoff is mainly due to reduced host mRNA levels ^47^.

Traditionally, IAV is thought to prioritize the translation of viral over host mRNAs ^48,49^, but more recent experimental and computational analyses challenge this view ^47,50^. We investigated this question by calculating protein synthesis efficiencies (i.e. the amount of protein made per mRNA). To this end, we divided iBAQ values by corresponding RPKM values (Figure 2F). Infection with both strains reduced host protein synthesis efficiencies compared to uninfected controls. Importantly, we did not observe preferential translation of viral transcripts. Instead, viral proteins were even less efficiently synthesized than host proteins in both strains. This suggests that mRNAs from human and avian Influenza virus strains access the translational machinery with comparable efficiency, which argues against the idea that modulation of translation efficiency affects species specificity.

### Dysregulated synthesis of viral proteins

Since the observed differences in host protein synthesis were surprisingly subtle we focused our attention to the dynamics of viral protein synthesis. Production of most viral proteins peaked in the 8-12 hour period (see Supplementary Figure 2). The kinetics such as the early production of NS1 and NP and delayed synthesis of M1 is consistent with classical radioactive pulse labeling experiments ^33^. We then used SILAC ratios of shared peptides (that is, peptides with sequence identity between both strains) to precisely compare the kinetics of viral protein synthesis (Figure 3A-B). We found that the avian strain produced higher amounts of all viral proteins at the beginning, confirming that the Mal virus successfully enters cells and initiates its gene expression program. Later on, during mid to late phases, the human Pan virus produced most proteins more abundantly than the avian strain. Note that NS1 and M2 are excluded in this analysis because no identical peptides were identified.

**Figure 3.**
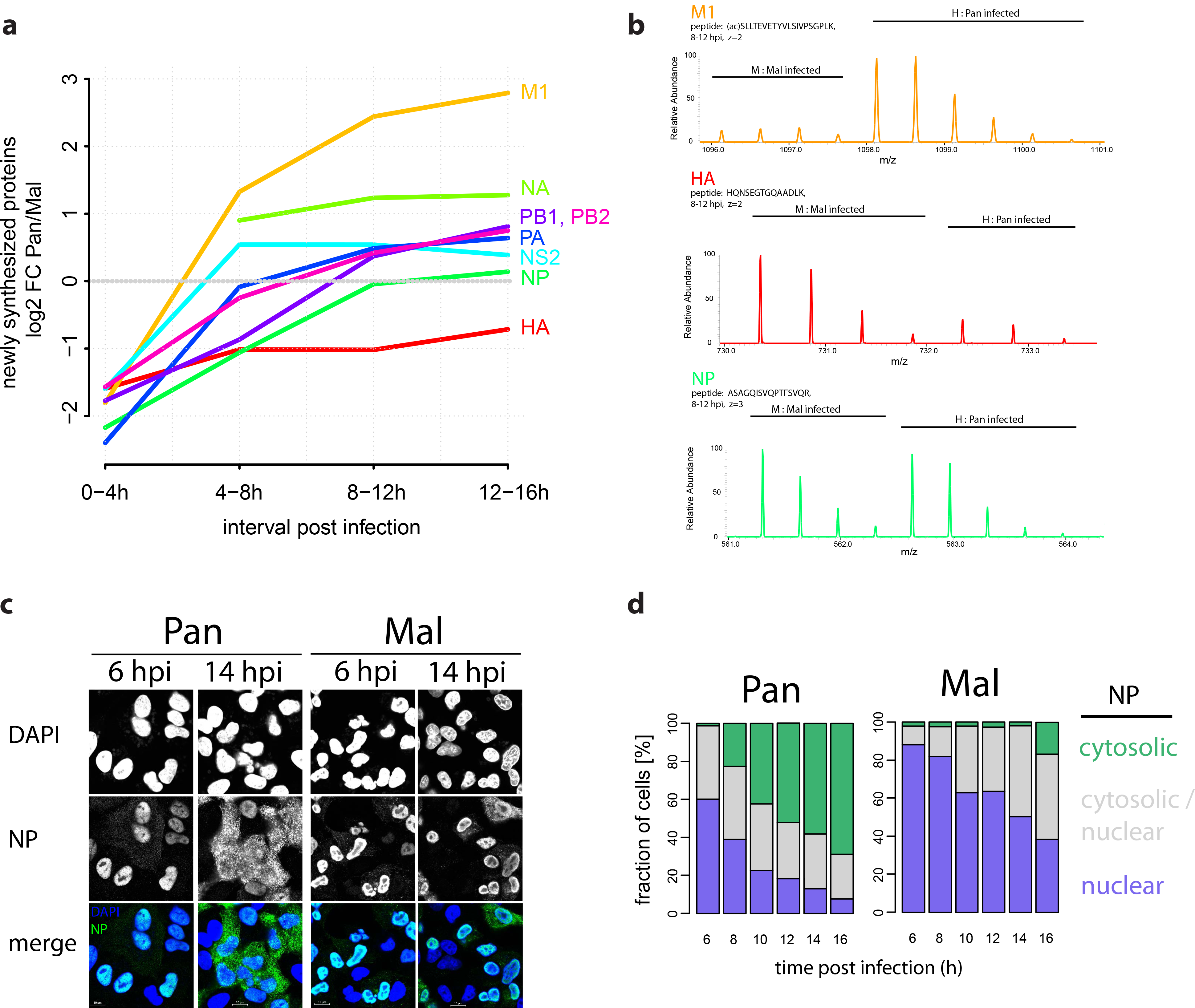
Viral protein synthesis is dysregulated during non-permissive infection. **a** SILAC protein synthesis profiles of human vs avian IAV proteins based on shared peptides. NS1 and M2 lack shared peptides. **b** MS1 spectra of individual precursor (M1 - top, HA - middle, NP - lower) peptides in M and H SILAC channels. **c** Cells were infected with the indicated viruses at MOI=1 (FFU/cell) and analyzed by immunofluorescence microscopy for NP trafficking. Nuclei were counterstained with DAPI. **d** For quantification, NP staining pattern was categorized as predominantly cytosolic, nuclear or both. At least 150 cells were counted per condition.

It is well-established that the RNA dependent RNA polymerase (RdRp) from avian-adapted IAV strains is less active in mammalian cells ^8,9^. Thus, we would have expected the production of all viral proteins in the bird-adapted strain to be reduced to a similar extent. In contrast, we observed striking differences in the synthesis of individual proteins: Hemagglutinin (HA) was more abundantly produced by the avian strain throughout infection. In contrast, neuraminidase (NA) and particularly matrix protein M1 were stronger produced by the human strain at later stages. These differences in the production of individual viral proteins cannot be explained by the global difference in RdRp activity between strains. Thus, the avian strain displays dysregulated protein production relative to its human counterpart (Figure 3A).

We focused our attention on the M1 protein since it showed the largest difference between both strains. The protein is highly conserved between Pan and Mal (~96% amino acid identity) and the most abundant protein in virions ^51^. Moreover, M1 is known to mediate export of the viral genome across the nuclear membrane - an essential step during permissive infection ^15,16^. Thus, accumulation of M1 at late stages of infection is required for the appearance of viral ribonucleoproteins (vRNPs) in the cytoplasm of infected cells. Interestingly, when investigating the subcellular distribution of the viral nucleoprotein (NP) by immunofluorescence microscopy, we observed efficient export during infection with the Pan strain (Figure 3C). In contrast, NP was inefficiently exported and accumulated in the nucleus upon Mal infection. These microscopy data is also corroborated by the increased interferon response induced by the Pan strain at later stages of infection (Figure 2C), which is stimulated by cytosolic viral RNA sensors ^52^. We conclude that non-permissive infection correlates with reduced M1 production and impaired nuclear export of NP.

### Non-permissive infection is characterized by increased M1 mRNA splicing

We next sought to investigate the mechanism for the impaired M1 production. To this end, we first quantified the levels of viral mRNAs from our RNA-seq data. In total, the avian virus produced ~2/3 of the mRNA of the human strain with the single largest difference observed for M1 (Figure 4A). The strain-specific differences in M1 mRNA levels were very similar to the observed differences in M1 protein production (Figure 4B). Hence, the impaired M1 protein production during non-permissive infection can largely be explained by reduced M1 mRNA levels.

**Figure 4.**
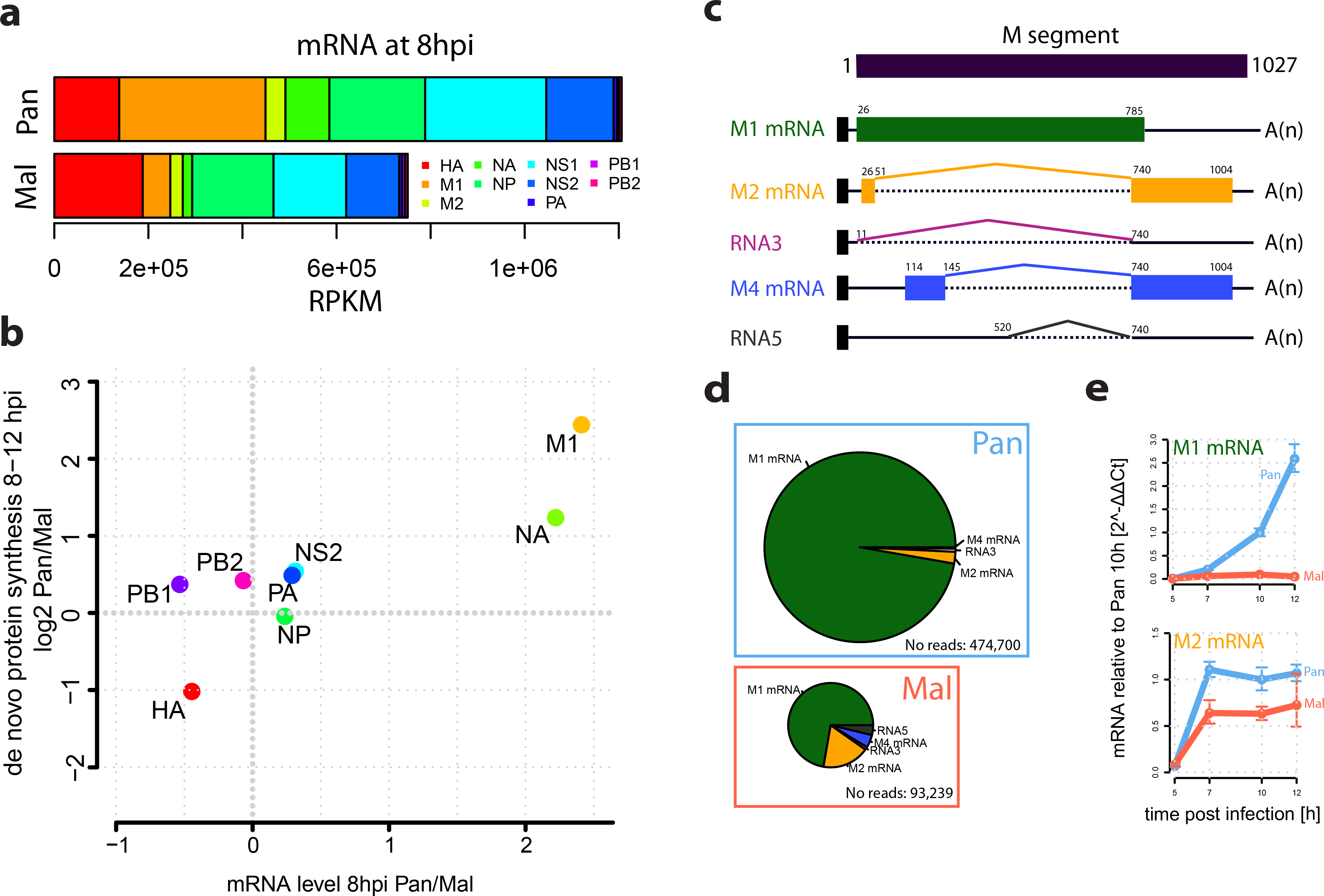
M1 splicing is markedly different in permissive versus non-permissive infection. **a** Normalized read counts (RPKM) for 10 viral mRNA from poly-A enriched samples from either Pan or Mal infected samples at 8 h p i. **b** Direct comparison of fold-changes for viral mRNAs/ proteins from RNA level and protein synthesis data. **c** Schematic depiction of M gene architecture. The M pre-mRNA can be spliced into the indicated isoforms. **d** Relative quantification of the different isoforms based on splice junction reads from RNAseq data for both strains at 8 h p i. The area of the pies reflects the absolute number of splice junction reads. **e** M1/M2 mRNA abundance kinetics based on qRT-PCR of cells infected with the indicated viruses at MOI=4 (FFU/cell).

M1 is encoded on segment 7 (that is, the M segment), which is the most conserved segment between Pan and Mal (~89 % nucleotide identity). The M1 protein is produced from a collinear transcript that can be alternatively spliced into three additional isoforms which all use a common 3’ splice site ^53,54^: the M2 mRNA, which encodes the ion channel M2 ^17^, RNA 3 which is not known to encode a peptide, and M4 mRNA that is proposed to be translated to an isoform of the M2 ion channel in certain strains ^55^. We investigated the relative proportion of these isoforms in the RNA-seq data via splice junction reads. We detected all known isoforms plus a novel transcript of the avian M segment, which we call RNA 5. This transcript results from splicing at 5’ donor GG site (pos 520/521) and the common 3’ acceptor site and contains an ORF in-frame with M1 with a missing internal region (Figure 4C).

While only a few percent of the M1 mRNA was alternatively spliced during permissive infection, ~1/3 was spliced upon infection with the avian strain (Figure 4D). Thus, the reduced level of M1 mRNA during non-permissive infection is at least partially due to increased splicing of the M1 mRNA to alternative isoforms. To validate these data we assessed the kinetics of M1 and M2 mRNA levels during infection via qRT-PCR (Figure 4E). This confirmed the reduced levels of M1 mRNA in the avian strain, especially at later stages. In contrast, M2 mRNA levels were overall similar throughout infection. We note that the comparable M2 mRNA level during non-permissive infection results from two opposing processes -- the increased splicing of the primary transcript to the M2 mRNA, and the global reduction in viral transcripts, which is probably due to the impaired polymerase activity ^8,9^. We conclude that M1 splicing is markedly different in permissive versus non-permissive infection.

### Difference in M1 mRNA splicing is determined by a cis regulatory element at the 3’ splice site

The differences in M1 mRNA splicing can be due to (i) cis-regulatory elements (that is, specific signals encoded in the M segment), (ii) trans-acting factors (that is, other viral or host factors that interact with M1 mRNA) or (iii) a combination of both. To assess whether cis regulatory elements are involved, we sought to investigate M1 splicing outside the context of infection. We therefore designed a splicing reporter system (Figure 5A-B). To this end, we cloned the coding region of the M segment (nt 29-1007) into a eukaryotic expression vector and fused it to an N-terminal Flag/HA tag. Importantly, this construct avoids the strong 5’ splice site of mRNA 3 ^56^ and enabled us to assess the relative levels of M1 to M2 proteins and mRNAs. When we transfected human A549 cells with these reporter constructs we found that M2 was produced to high levels with the construct containing the Mal M sequence but was barely detectable when the Pan M sequence was transfected (Figure 5C). Thus, our reporter system recapitulates splicing differences observed during infection. We conclude that cis-regulatory elements in the M segment cause excessive splicing of the avian variant.

**Figure 5.**
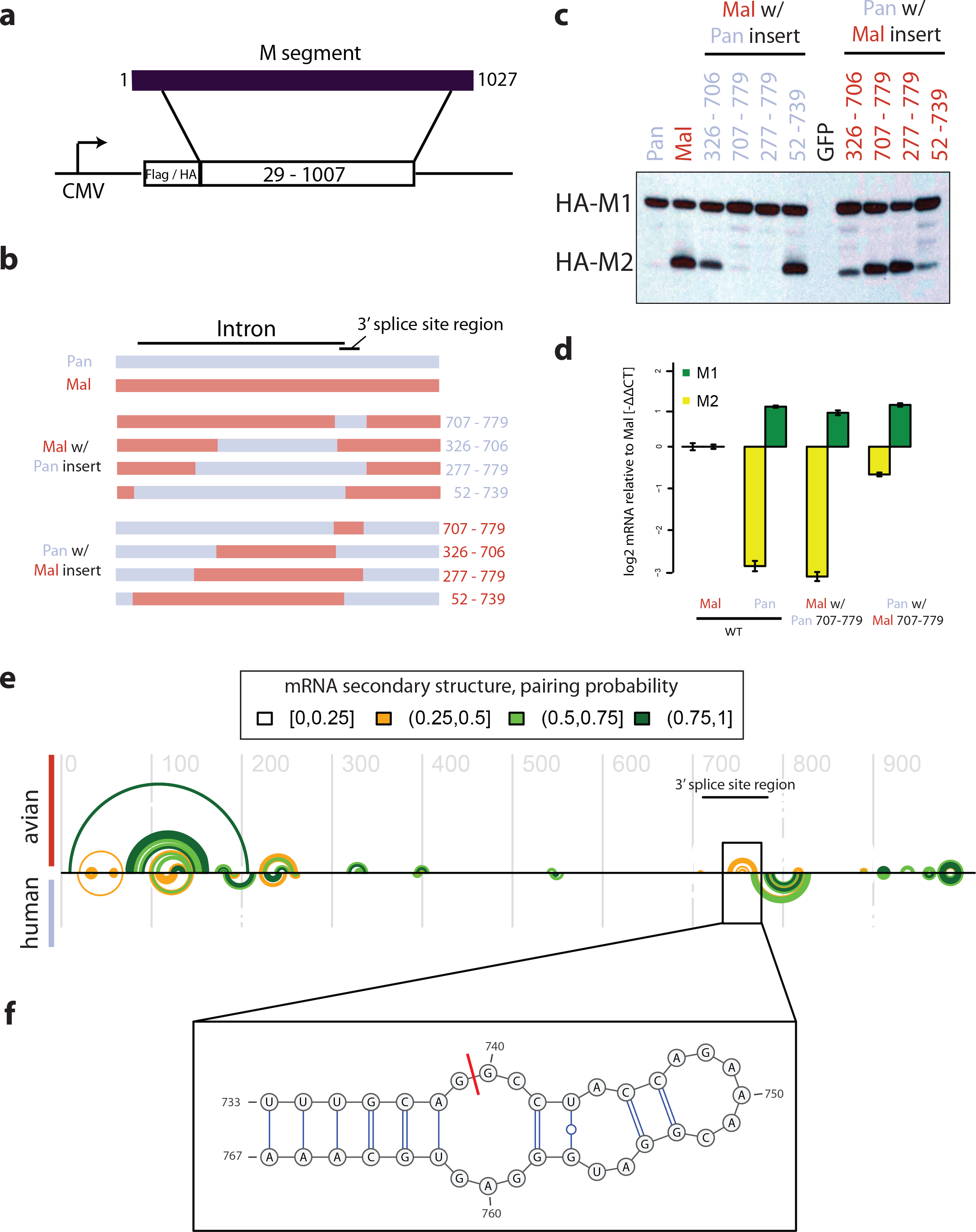
A cis-regulatory sequence element in the 3’ splice site controls M segment splicing efficiency. **(a)** Reporter system. The coding sequence of segment 7 was cloned into an eukaryotic expression vector with N-terminal Flag/HA tag. **(b)** Wildtype Pan and Mal sequences as well as several chimeric Pan/Mal constructs were cloned. A549 cells were transfected with the indicated reporter constructs, harvested and subjected to anti-HA immunoblotting **(c)** or qRT-PCR **(d). (e)** Evolutionary conserved RNA secondary structure along the M mRNA predicted for avian adapted (top) and human adapted (bottom) IAV strains. Each semi-circle corresponds to one base-pair involving the corresponding two alignment positions. Colour-coding of base-pairs according to their corresponding, estimated base-pairing probability. The 3’ splice site region is annotated with a bar. **(f)** The insert shows a detail of the predicted RNA secondary structure at the 3’ splice site, predicted for avian but not human sequences in this study. The splice position is indicated by a red bar.

To determine the sequence responsible for the strain-specific splicing we made chimeric reporter constructs (Figure 5B). When swapping the entire intron sequence of the M2 splice variant (nucleotides 52-739, corresponding to ~70% of the CDS), we did not observe major changes in the relative amount of M1 to M2. In contrast, integrating the human 3’ splice site region (nucleotides 707-779, 73 nucleotides) into the avian construct strongly impaired splicing down to the levels of the human wild type construct. Conversely, when we integrated the avian 3’ splice site region into the human construct we observed a strong increase in splicing, similar to the avian wild type construct (Figure 5C). To validate these results at the mRNA level we used qRT-PCR. Again, we found that the splice site region alone is sufficient to switch the species-specific splicing phenotype (Figure 5D). Interestingly, this region has been reported to contain an RNA secondary structure ^57^ and a binding site for the splicing factor SRSF1 ^58^. We conclude that a cis regulatory element in the splice site region determines the strain-specific splicing pattern.

### An RNA hairpin that spans the 3’ splice site is evolutionary conserved in avian but not in human-adapted M segments

We next wanted to assess whether our findings are also relevant for other human-or bird-adapted IAVs. Specifically, we sought to identify functionally relevant RNA secondary structures that have been conserved during evolution of avian-and human-adapted IAVs. To this end, we analysed multiple sequence alignments from hundreds of recent human and avian H3N2 isolates using the RNA structure prediction program RNA-Decoder ^59^(see Material and Methods). This program is capable of dis-entangling overlapping evolutionary constraints due to encoded amino acids and RNA structure features and has been shown to successfully identify evolutionarily conserved RNA structures overlapping protein-coding regions, e.g. in viral genomes such as hepatitis C and HIV ^59,60^. Importantly, RNA-Decoder captures evidence on conserved RNA structure based on the evolutionary signals encoded in the sequences of the input alignment. This is a key advantage over computational methods that identify RNA structures based on their thermodynamic stability *in vitro,* as these methods assume that the RNA has no interactions with other molecules (e.g. proteins and other RNAs) *in vivo*. Also, RNA-Decoder employs a probabilistic framework which is capable of estimating the reliability of its predictions.

The RNA secondary structure that is best supported by the evolutionary signals in the two multiple sequence alignments (the so-called maximum-likelihood structure) markedly differs between human and avian strains, particularly in the region around the 3′ splice site (Figure 5E): The avian region encodes a hairpin-like structure (Figure 5F) overlapping the 3′ splice site which is absent from the human-adapted sequences. This structure is similar to a hairpin reported by Moss et al. for four sequences ^57^ but differs in details (Supplementary Figure 3). Most importantly, the evolutionarily conserved structure reported here leaves the GC-motif immediately downstream of the AG consensus at the 3′ splice site unpaired, making it potentially more accessible to splicing. We conclude that the M segment of avian and human-adapted H3N2 isolates contain evolutionarily conserved RNA secondary structures that markedly differ in exactly the region that is critical for strain-specific splicing.

In addition to these computational analyses, we also wanted to test the relevance of our findings for other IAV isolates experimentally. The M segment of the seasonal Pan strain originates from the M segment of the A/Brevig Mission/1/1918 (p1918) virus, which is at the evolutionary root of human strains and caused the 1918 “Spanish flu” pandemic ^61^. Therefore, we cloned the M segment of p1918 into our reporter vector (Supplementary Figure 4). Again, we observed inefficient splicing of the p1918 M gene, consistent with our data for the Pan strain and previous reports ^62^. Moreover, integration the Mal 3’ splice site region into the p1918 gene increased splicing. Thus, inefficient splicing of the M gene in human-adapted IAVs occurs in a seasonal (Pan) and a pandemic (p1918) strain.

### The 3’ splice site is a host range determinant

The experiments with reporter constructs described above are advantageous because they allow us to study the impact of M segment sequence features in isolation. Nevertheless, it is also important to assess the relevance of these findings during infection. We therefore mutated eight nucleotides in the splice site region of the Pan wild type strain to the corresponding nucleotides in the Mal strain using reverse genetics (“Pan-Av” for a Pan strain with an avian splice site region, see Figure 6A and Supplementary Table S3).

**Figure 6.**
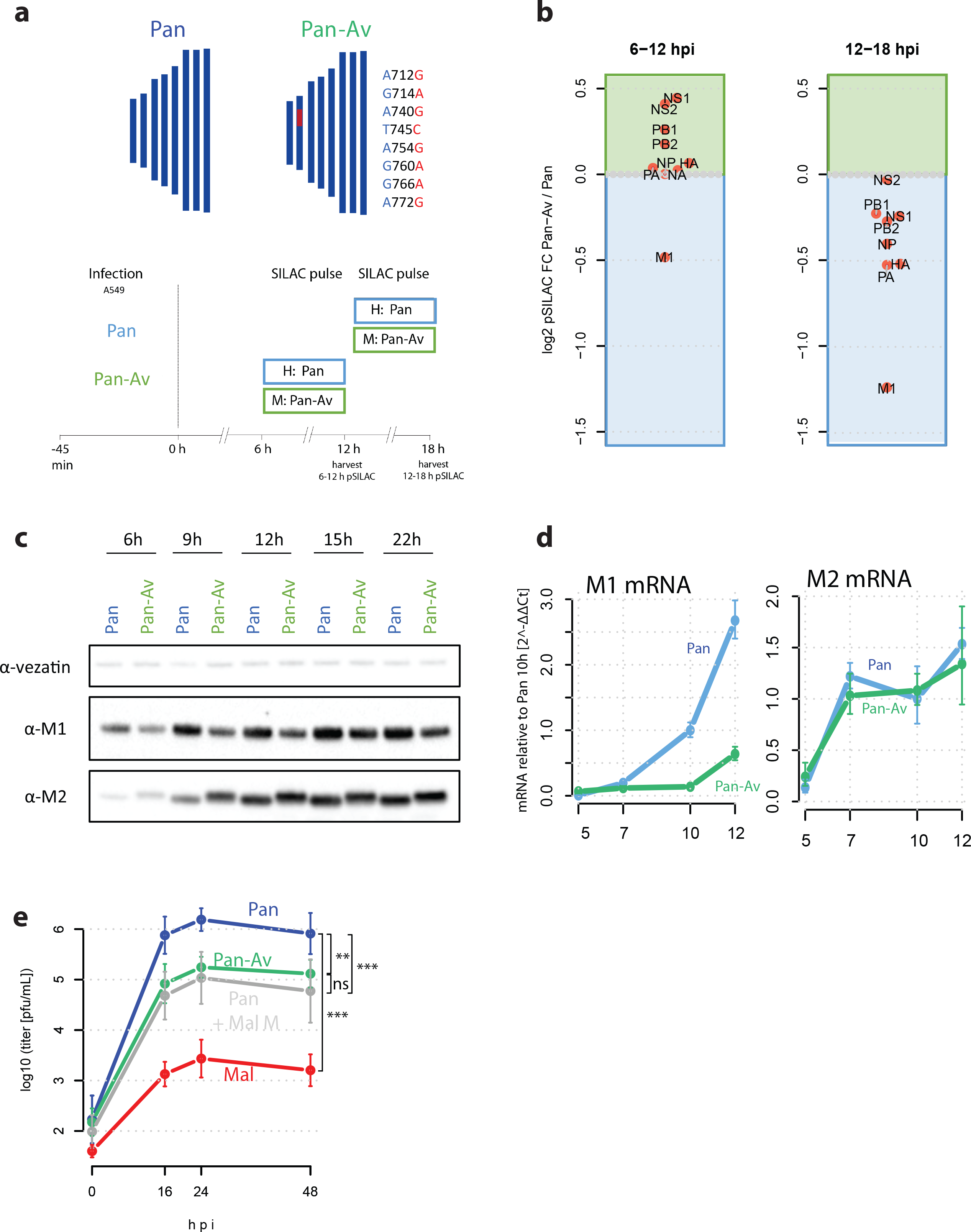
The 3’ splice site is a host range determinant. **a**, top 8 nucleotide polymorphisms of Mal virus at the segment 7 3’ splice site were integrated into wildtype Pan virus generating “Pan-Av” mutant. **a**, bottom Design of pulsed SILAC experiment. A549 cells were infected with wildtype or mutant virus at an MOI of 4 (FFU/cell) and viral gene expression was assessed by pulse labeling during two time intervals. **b** SILAC-based quantification of viral protein expression comparatively for Pan and Pan-Av virus. Kinetics of M1/M2 expression was detected by immunoblotting **c** or detected by qRT-PCR **d**. **e** Multicycle replication curve of the indicated viruses at an MOI of 0.05 (FFU/cell) on A549 cells. Means and standard deviations of biological triplicates are shown along with significance estimates based on paired t-tests for the 16-48 h time points (ns: non-significant, **: p<0.01, ***: p<0.001).

We first compared the kinetics of viral protein synthesis upon infection of A549 cells with both strains using pSILAC ^35^. M1 synthesis was selectively impaired during Pan-Av infection during both the 6-12 and 12-18 hpi time intervals. At later stages, the Pan-Av strain also showed impaired production of other essential viral proteins (Figure 6B), suggesting that viral replication is also impaired. Next, we quantified M1 and M2 protein (Figure 6C) and mRNA levels (Figure 6D). The Pan-Av strain displayed decreased M1 protein and mRNA levels, mimicking the behaviour of the Mal strain (compare also Figure 4).

To assess the impact of M segment splicing on IAV replication in human cells, we assessed the growth characteristics of the different viruses (Figure 6E). As expected, the Pan strain reached ~1,000 fold higher titers than the Mal strain. Exchanging the entire M segment of the Pan strain with the M segment of the Mal strain (Pan + Mal M) reduced titers about 10-fold. Importantly, a similar ~10 fold attenuation was also seen in the Pan-Av strain that only differs from the Pan strain by 8 nucleotides. We conclude that the 3’ splice site of the IAV M segment is indeed an important host range determinant.

## DISCUSSION

Advances in high-throughput sequencing have provided insights into the extraordinary diversity of viruses and their genomic determinants of host adaptation. However, the mechanism how these adaptive mutations enable replication in a given host is less understood. Our proteomic pulse labelling data allowed us to take an unbiased look at protein synthesis upon permissive and non-permissive infection. We found that the synthesis profiles of host cell proteins were remarkably similar. Hence, the outcome of infection does not appear to depend on a specific host response. In contrast, we observed striking differences in the synthesis profiles of viral proteins. Particularly, the matrix protein M1 was inefficiently produced during non-permissive infection. Our follow-up experiments showed that this depends -- at least partially -- on excessive splicing of the avian M1 mRNA to alternative transcripts. Systematic computational analysis of the RNA structure of the M segment revealed characteristic and evolutionarily conserved differences in the splice site regions between human and bird-adapted strains. Exchanging eight nucleotides in the 3’ splice site region from the human-adapted strain to corresponding sequences in the bird-adapted strain markedly impaired replication. Thus, our proteomic analysis of IAV infection identifies M segment splicing as a host range determinant. A hypothetical model for the influence of M segment splicing on the IAV host range is presented in Figure 7.

**Figure 7.**
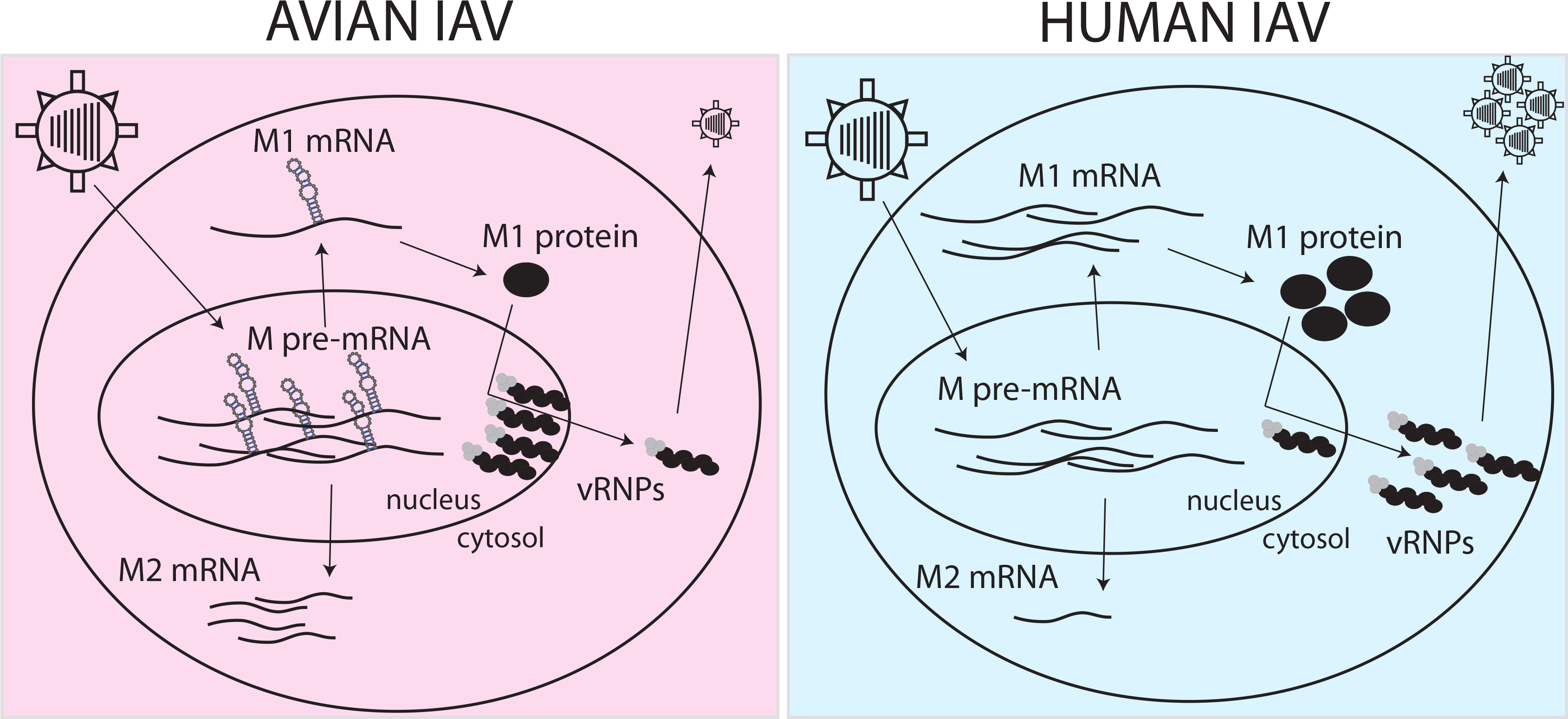
Hypothetical model for the role of M segment splicing for IAV host range. A cis-regulatory secondary structure element (indicated hairpins) in avian but not human IAV M pre-mRNAs facilitates splicing. This leads to the underproduction of M1 mRNA and protein in human cells infected with avian-adapted IAVs. The poor availability of the M1 protein may contribute to an impaired nuclear export of viral ribonucleoproteins (vRNPs).

The cell culture-based infection model and splicing reporter system employed here are advantageous because they enable experiments under well-controlled conditions. Having said this, it is important to keep in mind that these model systems do not represent the full complexity of IAV infections *in vivo*. Also, the Pan and Mal strain employed here do not represent the full diversity of human-and bird-adapted IAVs. However, our computational analyses show that differences in RNA secondary structure of the 3’ splice site are widely conserved in human-and bird-adapted IAVs. Moreover, recent results from the Steel lab show that avian M segments restrict growth and transmission of mammalian-adapted IAV strains in a guinea pig model (John Steel, Emory University, personal communication). Together, these findings strongly suggest that our results are also relevant outside the specific experimental model system employed here.

Our global assessment of protein *de novo* synthesis upon infection revealed a global reduction in overall protein output during infection with both strains, probably reflecting a global stress response. Also, we observed the well-known shutoff of host protein synthesis^44,46–48^. Specific classes such as interferon-related, ribosomal and mitochondrial proteins escaped the shutoff. We observed that the amount of protein synthesis upon infection primarily depends on mRNA levels. Thus, altered translation does not play a major role for the host shut-off, consistent with recent findings ^47^. Surprisingly, we also found that viral transcripts were not more efficiently translated than host transcripts. Instead, their translation efficiency (that is, the amount of protein made per mRNA) was even lower than for host proteins. This contrasts with early studies based on reporter systems ^48^ but corroborates recent ribosome profiling data ^47^. Our finding is also consistent with the fact that the codon usage of IAV genes is not optimized to reflect the codon usage of the host ^50^. It is also interesting that the translation efficiency of the bird-and the human-adapted strains was similarly poor. Thus, adaptation towards high translational efficiency does not seem to be required for crossing the species barrier.

Our unbiased proteomic analysis indicates that the differences between permissive and non-permissive infection depend on differences in viral rather than host protein synthesis. Hence, the orchestrated synthesis of the viral proteome appears to be critically important for permissive infection. This supports the emerging view that modulation of viral protein synthesis underpins host adaptation ^63^. Specifically, we find that the strain-specific differences in M1 protein synthesis critically depend on a conserved cis-regulatory element, which controls M-segment mRNA splicing. M1 is particularly important for the nuclear export of the viral genome to the cytoplasm ^15,16^. Consistently, we observed that the genome of the bird-adapted strain was inefficiently exported (Figure 3C).

We found that exchanging only eight nucleotides of the human-adapted M segment to the bird-adapted sequences markedly impaired viral replication. Hence, the cis-regulatory element described here plays an important role for host adaptation. However, it is critical to also emphasize that this is not the only relevant factor for IAV host range. For example, despite the overall similar host response, we and others have previously described host factors affecting human and avian virus infections ^20–23,25,32^. It is also well-established that the RdRp of avian-adapted strains is less active in human cells ^8,9^. Moreover, differences in the binding specificity of viral hemagglutinins (HA) are known to play an important role for host adaptation ^10^. Lastly, M-segment splicing does not only depend on cis-regulatory elements but also on trans-acting factors, such as NS1, RdRp, NS1-BP or HNRNPK ^56,64–66^. Indeed, while M1 production was clearly impaired in our mutant strain (Figure 6B), the wild-type bird-adapted strain produced even less (Figure 3A). It is therefore important to interpret our findings in the broader context of viral and host factors that jointly determine the success of IAV replication.

We are living in a pandemic era of IAV infections that began at around 1918 ^67^. At this time, a virus of ultimately avian origin acquired the ability to spread among humans and later on contributed its genetic material to other pandemic viruses until present. The M gene of this p1918 virus is in some regions similar to bird-adapted sequences but shows important signatures of mammalian adaptation, especially at the 3’ splice site ^68^. Our results suggest that mammalian adaptation at the 3’ splice site was linked to modulating M-gene splicing, which may have been relevant for the emergence of the 1918 pandemics in humans.

## ACKNOWLEDGMENTS

This work was supported by the German Ministry of Education and Research (Virosign grant 0316180B). We would like to acknowledge the excellent technical assistance of Gudrun Heins. The authors thank Katrina Meyer for providing the pDEST26-Flag/HA plasmid and Koshi Imami for setting up the nLC system using monolithic columns.

## AUTHOR CONTRIBUTIONS

B.B. performed most of the experiments and data analyses. K.E. established the AHA pulse labeling technology and generated pAHA SILAC data together with B.B‥ A.S. and K.P. performed the infection experiments for proteomic analysis supervised by T.W‥ X.W. and J.H. generated and analyzed RNA-seq data supervised by W.C‥ I.H. performed qRT-PCR analysis supervised by L.W. and B.B‥ M.H., L.W. and B.V. contributed to cloning and transfection of reporter constructs. I.M.M. performed the RNA structure analyses. T.W. and M.S. conceived and supervised the study. B.B., L.W., T.W. and M.S. interpreted the data. B.B. and M.S. wrote the manuscript with input from all authors.

---

**Supplementary Figure 1 (related to Figure 1).**
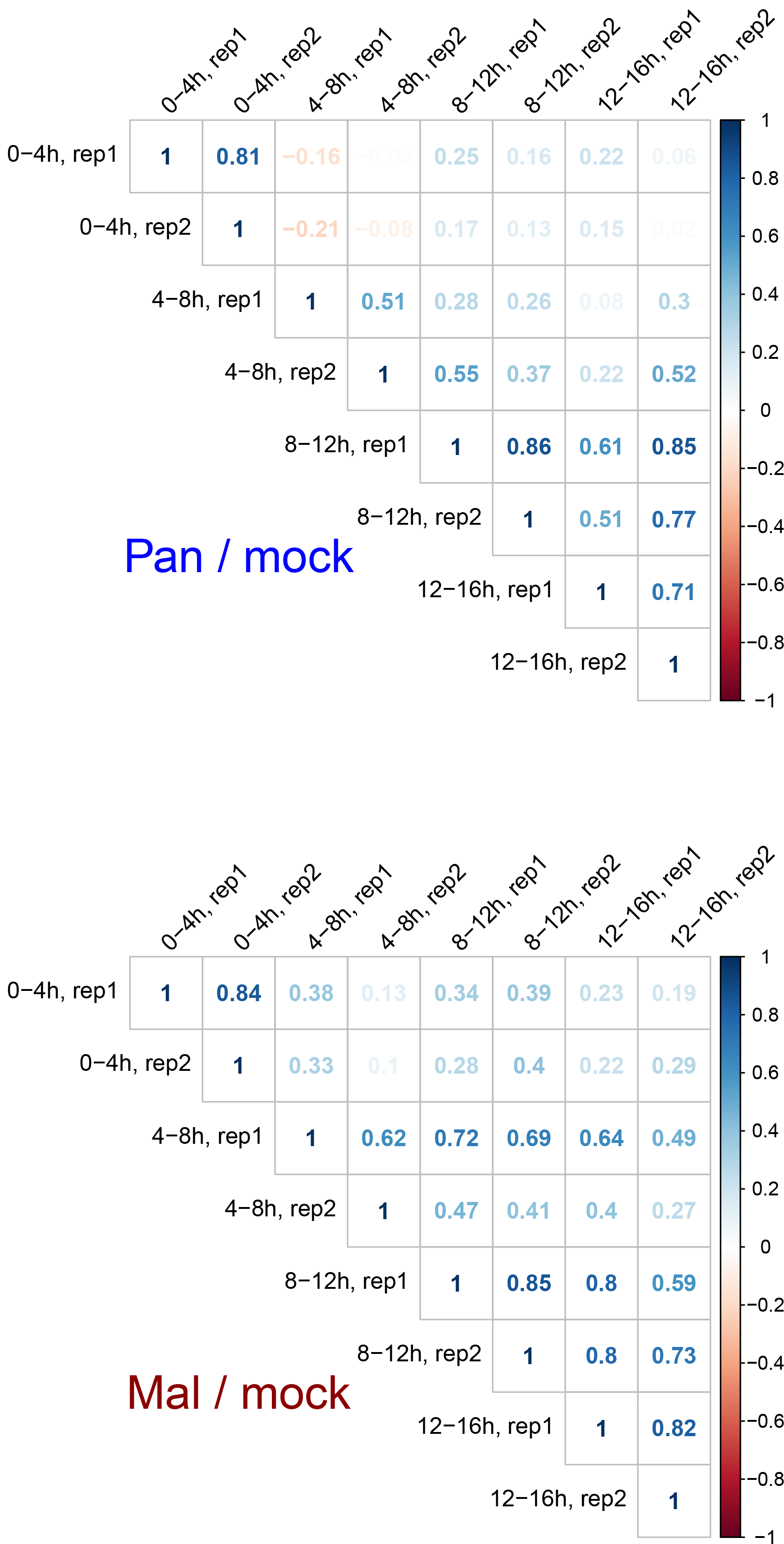
Data reproducibility. Spearman’s correlation coefficients of biological (label-swap) replicates for SILAC ratios Pan/mock (top) and Mal/mock (bottom).

**Supplementary Figure 2 (related to Figure 3).**
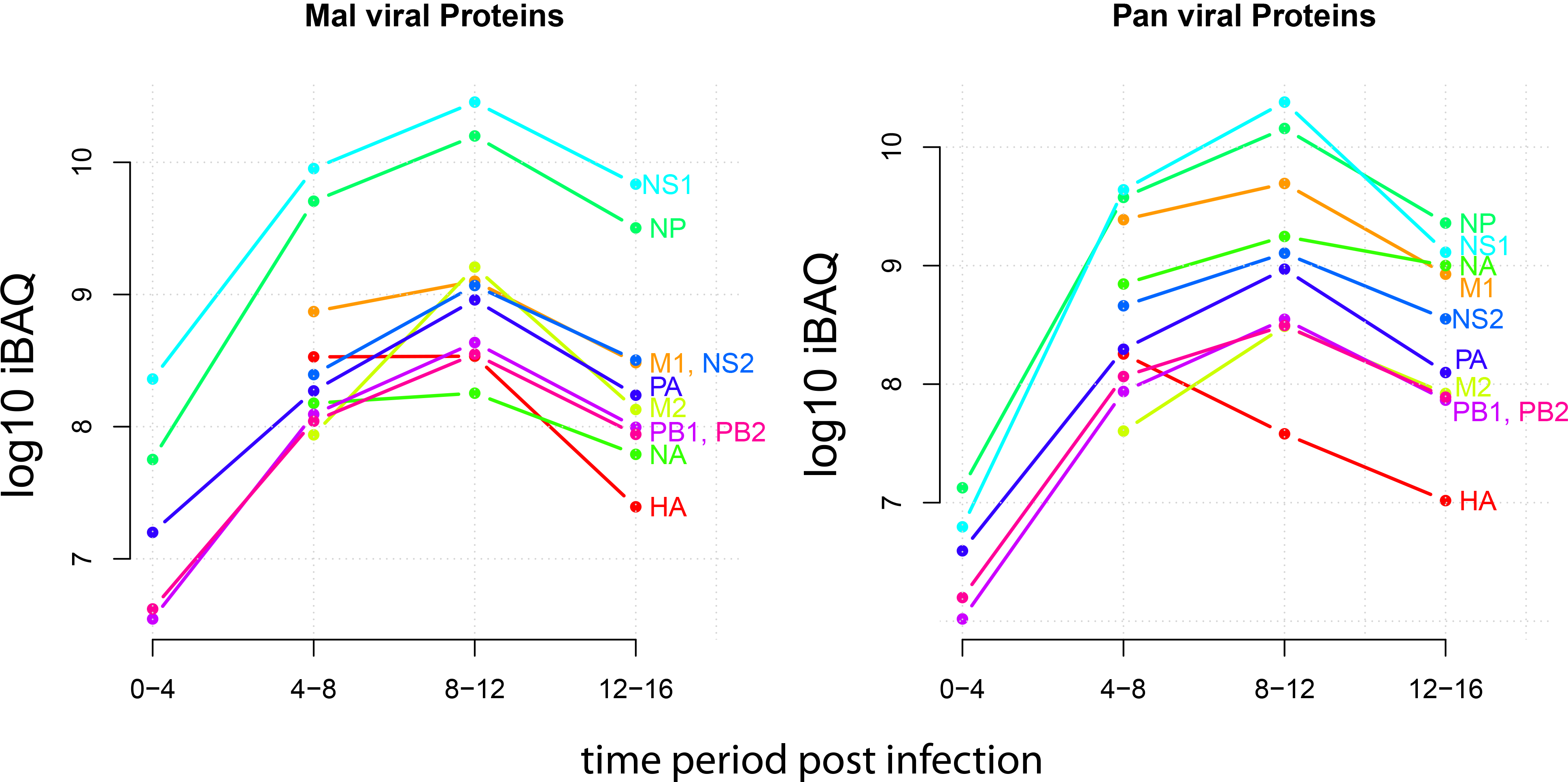
Viral protein expression kinetics. IBAQ-based absolute quantification of protein synthesis across infection for the indicated 10 viral proteins of strain Mal (left) or strain Pan (right).

**Supplementary Figure 3 (related to Figure 5).**
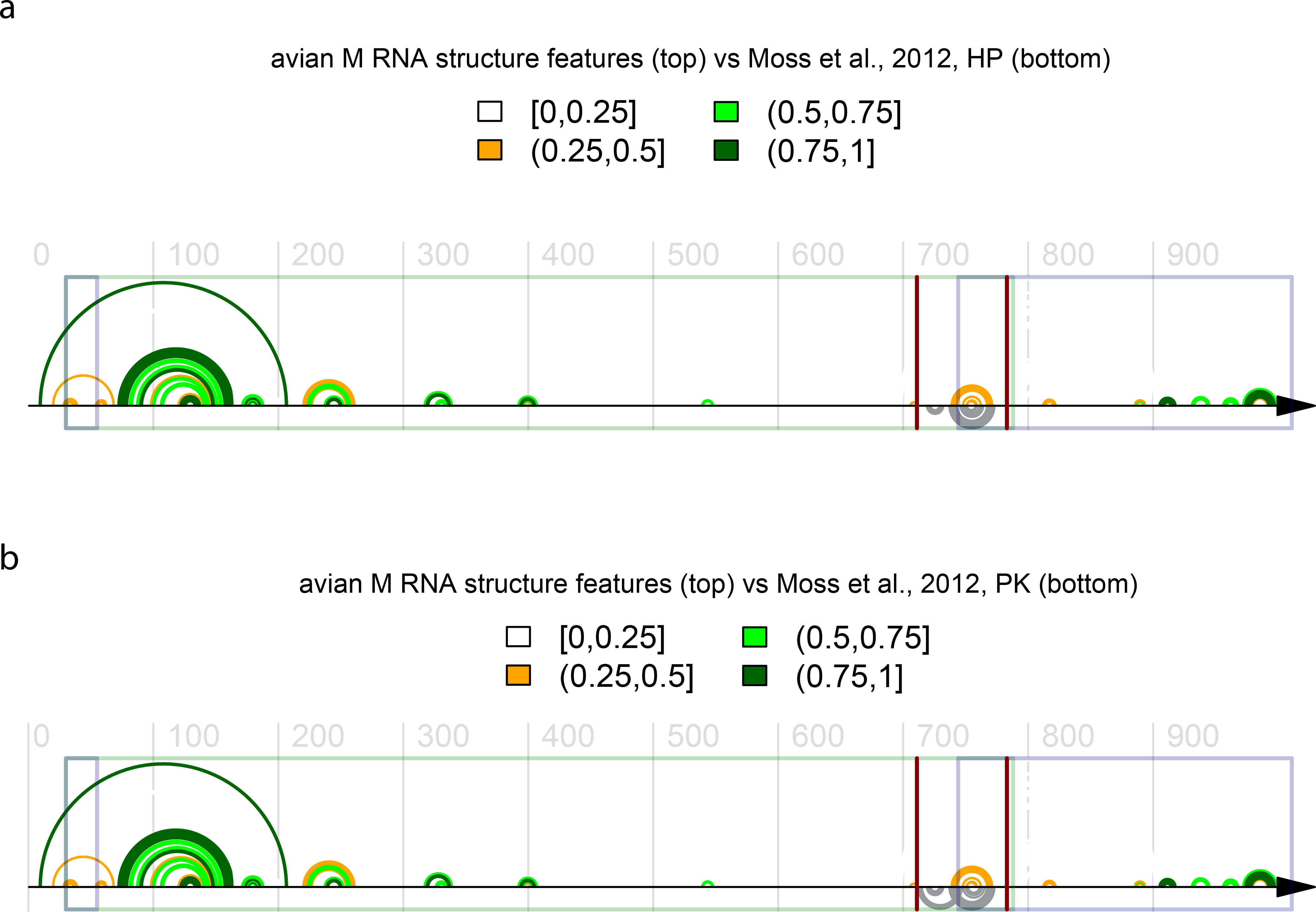
RNA structure comparison to published data. Agreement between the RNA secondary structure predicted by RNA-Decoder for the M mRNA for the avian-adapted influenza strains and the RNA structure features predicted by Moss et al., *PloS One*, 2012. Agreement with the hairpin (HP) in **a** and with the pseudoknot (PK) in **b.** Colour-coding of base-pairs according to the corresponding, estimated base-pairing probabilities, see also legend of Figure 5. The single exon containing the contiguous open-reading frame (ORF) of the M1 splice variants is indicated by a green box, the two exons corresponding to the M2 splice variant are shown as two blue boxes. Note that there are two regions where the ORFs of the splice variants overlap. The first one corresponds to the first exon of the M2 splice variant where the two ORFs are in sync. The second one corresponds to the region of overlap between the 3′ end of the long M1 ORF and the 5′ start of the second exon of the M2 splice variant. In that region, the two ORFs are out of sync, implying a particularly strong constraint due to the two, intertwined amino-acid contexts. The two vertical bars indicate the start and end position of the region that was used for chimeric constructs around the 3′ splice site.

**Supplementary Figure 4.**
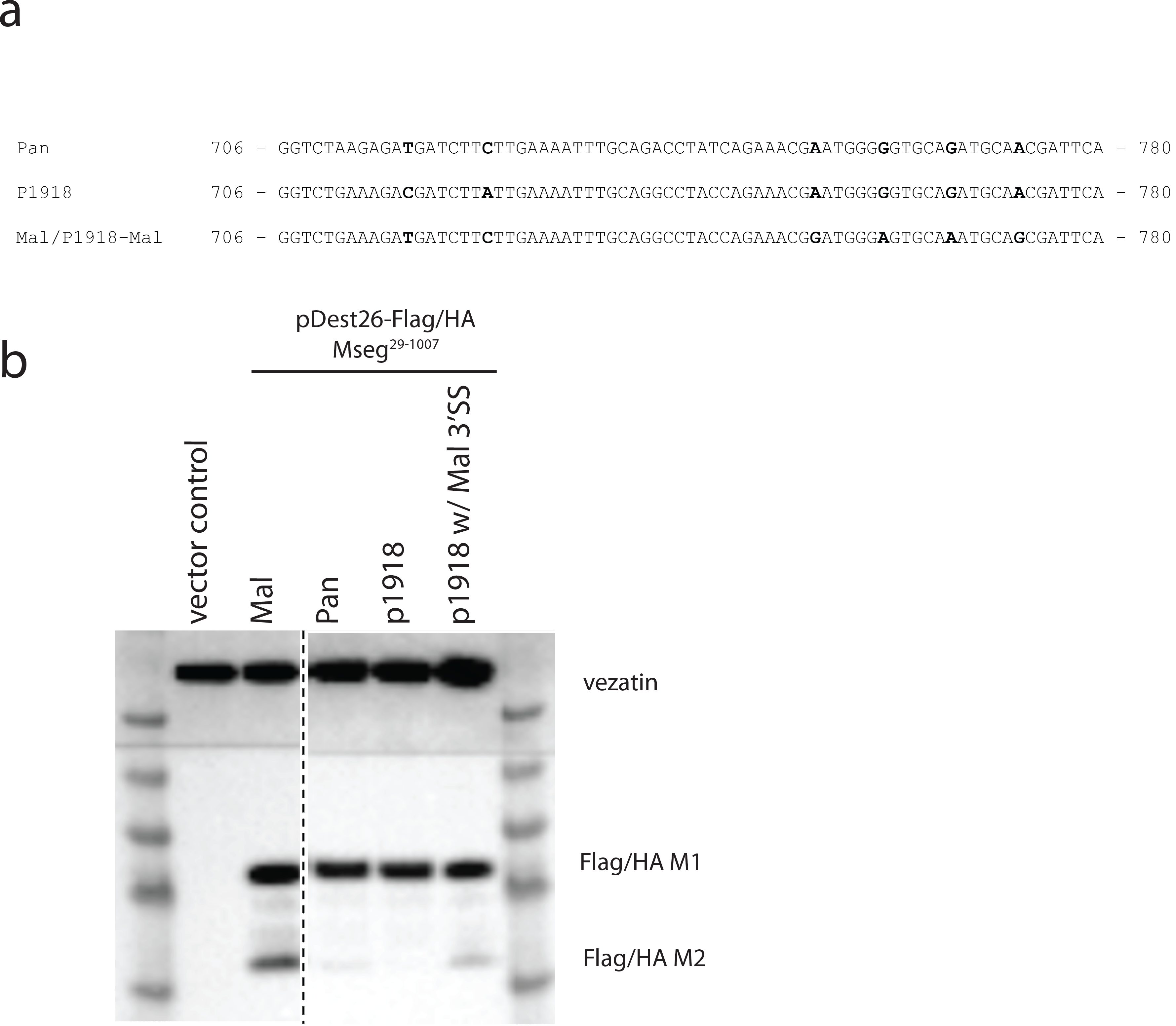
Pandemic 1918 M gene is spliced poorly (related to Figure 5) **a** Multiple sequence alignment of the nucleotide sequences of strain Pan, Mal and p1918 in the M segment 3’ splice site region. A mutant reporter construct was created that comprises the Mal 3’ splice site region in the p1918 M segment backbone (p1918-Mal). **b** A549 cells were transfected with the indicated expression constructs and M1 / M2 expression was assessed by immunoblotting against HA-antigen.

## MATERIAL AND METHODS

### Cells and Viruses

A549 (human) were grown in Dulbecco modified Eagle medium (DMEM) supplemented with 10% (v/v) fetal bovine serum, 2 mM L-glutamine, and antibiotics. MDCK type II cells were grown in minimal essential medium (MEM) supplemented with 2mM L-glutamine. All cells were maintained at 37 °C and 5% CO2. Stocks of the avian influenza viruse A/Mallard/439/2004 (H3N2) (Mal) (GISAID accession numbers EPI859640-EPI859647) were grown in the allantoic cavities of 10-day-old embryonated chicken eggs for 2 days at 37°C. A/Panama/2007/1999 (H3N2) (Pan) (NCBI accession numbers: <u>DQ487333-DQ487340</u>), Pan + Mal M reassortant and Pan-Av mutant virus were grown in MDCKII cells. Virus stocks were titrated on MDCK type II cells by measuring plaque forming units (PFU) or fluorescence forming units (FFU). For the latter, infected cells were infected with different dilutions of allantoic fluid for 5 h. Then cells were harvested by trypsinization, fixed and permeabilized by incubation in 75% ethanol for at least 12 h at 4°C and stained with specific antibody against NP antigen (clone AA5H, Serotec). An Alexa Fluor 488-conjugated goat anti-mouse IgG antibody (Invitrogen) was used as secondary reagent. Cells were analyzed on a FACSCanto II flow cytometer (BD Biosciences) using FACSDiva software package.

### Cloning and mutagenesis

Gene fragments containing the coding sequence (nt 29-1007) of segment 7 of A/BrevigMission/1/1918 (GenBank accession: AY130766) and of A/BrevigMission/1/1918 with the following point mutations: C718T, A725C, A754G, G760A, G766A, A772G (p1918-Mal) fused to attB1 and attB2 sites were ordered as synthetic double-stranded DNA fragments from Integrated DNA Technologies. The coding sequences of Pan and Mal M segments (nt 29-1007) were amplified from cDNA and fused to attB1 and attB2 sites by PCR. Cloning was done using Gateway technology (Invitrogen) according to the manufacturer’s protocol. Entry clones were generated from pDONR221 vector, expression clones from pDEST26-Flag/HA destination vector. Pan/Mal chimeric constructs were generated by replacing appropriate restriction fragments of wild type pDEST26-Flag/HA-Pan and pDEST26-Flag/HA-Mal respectively with the corresponding synthetic chimeric DNA inserts purchased as gBlocks from Integrated DNA Technologies.

A cDNA copy of the M segment of A/Mallard/439/2004 was amplified by RT-PCR followed by insertion into pHW2000 resulting in pHW2000-Mal M. Also, the pHW2000-Pan M plasmid was avianized by introduction of eight point mutations into the M1 reading frame within pHW2000-M (A712G, G714A, A740G, T745C, A754G, G760A, G766A, A772G) using the Gibson Assembly cloning kit (NEB) (pHW2000-Pan-AV). The pDEST26-Flag/HA-Pan w/ Mal 707-779 and the wild type pHW2000-Pan M served as template for PCR. All constructs were confirmed by cycle sequencing.

### Reverse genetics

Recombinant influenza A viruses derived from the A/Panama/2007/99 backbone were generated using an eight plasmid system for this strain based upon pHW2000 by transfection of human HEK293T cells followed by passage on MDCK II cells as described elsewhere ^69^. For the rescue of the Pan-Mal M reassortant virus, we used pHW2000-Mal M, whereas the Pan-AV virus was generated with pHW2000-Pan-AV together with seven plasmids encoding the other segments of human A/Panama/2007/99 virus.

### AHA-SILAC

A549 cells were fully labeled in SILAC DMEM (PAA) supplemented with glutamine, 10 % FBS (Life Technologies) and 2 mM L-glutamine, antibiotics and with either heavy (R10K8, “SILAC - H”), medium (R6K4, “SILAC - M”) or light (R0K0, “SILAC - L”) arginine and lysine (Cambridge Isotope Laboratories). Cells were cultured in SILAC - L/M/H medium for at least 6 passages. 10 cm dishes of confluent light labeled cells were mock-infected, while heavy and medium labeled cells were infected with either Pan or Mal strain at an MOI of 3 (PFU). Virus was allowed to attach to the cells for 45 min on ice. Cells were washed with pre-warmed PBS before infection medium was added (SILAC DMEM containing the respective SILAC AA, 0.2 % BSA, 2 mM glutamine, antibiotics). Prior to pulse labeling cells were washed with pre-warmed PBS. Methionine depleted infection medium additionally containing 100 μM L-Azidohomoalanine (Anaspec) was added for different 4 h intervals to the cells. Cells were washed in PBS, scraped from the dish and frozen until further sample processing. Lysis and enrichment for newly synthesized proteins was done using Click-It protein enrichment kit (Invitrogen), with slight modifications: 283 μl of urea lysis buffer was used per label, cell debris was removed before SILAC label were mixed. 10% of sample were directly subjected to Wessel-Flügge precipitation ^70^ and served as the input, 90% were used for enrichment of newly synthesized proteins as previously described ^71^. Enriched proteins were reduced and alkylated as indicated in the manufacturer’s instruction. Beads were then washed sequentially (each 5x) in SDS wash buffer (supplied with the kit), 8 M urea in 0.1 M Tris/HCl (pH 8.0), 80 % acetonitrile in 0.1 M Tris/HCl pH 8.0 and 5 % acetonitrile in 50 mM ammonium bicarbonate. Proteins were then digested in 5 % acetonitrile / 50 mM ammonium bicarbonate overnight using trypsin (Promega). Peptides were then acidified, desalted and either directly measured on a nano LC-MS/MSset-up (see below) or subjected before to isoelectric focusing using an OFFGEL-fractionator.

Input samples were reduced by adding DTT to a final concentration of 0.1 M and incubation for 5 min at 95 °C. Sulfhydryl groups were alkylated by adding iodoacetamide to a final concentration of 0.25 M and incubation for 20 min in the dark at room temperature. Proteins were precipitated according to Wessel and Fluegge^70^, resuspended in 6 M urea / 2 M thiourea and digested into peptides with C-terminal lysine or arginine using Lys-C (3 h) and Trypsin (overnight, diluted 4× with 50 mm ABC). Enzyme activity was quenched by acidification of the samples with trifluoroacetic acid. The peptides were desalted with C18 Stage Tips ^72^ prior to nanoLC-MS/MS analysis.

### pSILAC

Cells were adapted to SILAC light medium one day before the experiment and infected as described above using an MOI of 4 (FFU/cell). Prior to the pulse period, cells were maintained in PBS supplemented with Ca^2+^/Mg^2+^ and 0.2% BSA for 30 min. Then cells were pulse labeled with SILAC M or SILAC H medium for 6 h intervals, harvested and combined. Lysis was carried out in 125 mM NaCl, 0.1% SDS, 1% NP-40, 5% glycerol, 50 mM Tris-HCl, pH 7.4 for 1 h on a rotating wheel with subsequent centrifugation. The supernatant was precipitated according to Wessel and Flügge ^70^ and precipitated proteins were subjected to in-solution digest. Proteins were denatured in 2 M urea / 6 M thiourea, reduced, alkylated and digested using LysC (3 h at 20°C). Then, the digest solution was diluted 4x with 50 mM ammonium bicarbonate buffer and incubated with trypsin (Promega) for 16 h at 20°C. Afterwards, samples were acidified and subjected to stage tip purification as described previously ^72^.

### Mass Spectrometry

Peptides from input and AHA-enriched samples were separated on a monolithic silica capillary column (MonoCap C18 High Resolution 2000, GL Sciences), 0.1 mm internal diameter × 2000 mm length, at a flow rate of 300 nL/min with a 5 to 45% acetonitrile gradient on an EASY-nLC II system (Thermo Fisher Scientific) with 480 min gradient (Unfractionated AHA-enriched samples) or on a EASY-nLC HPLC (Thermo Fisher) system by 2 or 4 h gradients with a 250 nl/min flow rate on a 15 cm column with an inner diameter of 75 μm packed in house with ReproSil-Pur C18-AQ material (Dr. Maisch, GmbH). Peptides were ionized using an ESI source on a Q-Exactive, Q-Exactive Plus or a LTQ Orbitrap Velos MS (all Thermo Fisher) in data dependent mode. Q-Exactive and Q-Exactive Plus mass spectrometers were operated in the data dependent mode with a full scan in the Orbitrap followed by top 10 MS/MS scans using higher-energy collision dissociation. The full scans were performed with in a m/z range of 300 - 1,700, a resolution of 70,000, a target value of 3 × 10^6^ ions and a maximum injection time of 20 ms. The MS/MS scans were performed with a 17,500 resolution, a 1 × 10^6^ target value and a 60 ms maximum injection time. The LTQ Orbitrap Velos instrument was operated in data dependent CID top 20 mode. Full scans were performed in m/z range 300-1,700 with a resolution of 60,000 and a target value of 10^6^. MS/MS scans were performed with an isolation window of 2 m/z and a target value of 3,000.

Peptides from pSILAC samples were separated by 4 h gradients and ionized with ESI source and analyzed on Q-Exactive HF-X instrument (Thermo Fisher) in data dependent mode. The full scans were performed with a resolution of 60,000, a target value of 3 × 10^6^ ions and a maximum injection time of 10 ms. The MS/MS scans were performed with a 15,000 resolution, a 1 × 10^5^ target value and a 22 ms maximum injection time.

### Data Analysis

Raw files for AHA-SILAC were analysed with MaxQuant ^73^ software version 1.6.0.1 Default settings were kept except that ‘requantify’ option was turned on. Label-free quantification via iBAQ calculation was enabled. Lys4/Arg6 and Lys8/Arg10 were set as labels and oxidation of methionines, n-terminal acetylation and deamidation of asparagine and glutamine residues were defined as variable modifications. The in silico digests of the human Uniprot database (downloaded January 2018), the protein sequences of twelve Pan and Mal Influenza virus proteins and a database containing common contaminants were done with Trypsin/P. The false discovery rate was set to 1% at both the peptide and protein level and was assessed by in parallel searching a database containing the reverted sequences from the Uniprot database. The resulting text files were filtered to exclude reverse database hits, potential contaminants and proteins only identified by site (that is protein identifications that are only explained by a modified peptide). Plotting and statistics were done using R and figures were compiled in Illustrator (Adobe). Raw files for pSILAC were analysed as described above, except that MaxQuant software version 1.5.2.8 was used and requantify option was set to off.

### Proteomic data processing

Two MaxQuant output files were used: proteinGroups.txt and evidence.txt. iBAQ values from infected samples were extracted from proteinGroups.txt. iBAQ values were first normalized by scaling to the iBAQ protein median across all mock infected samples. This assumes that there are no differences in overall protein synthesis for different mock-infected samples. The iBAQ values were averaged for the corresponding label-swap replicates and proteins were categorized as host or viral. For estimating the newly synthesized protein mass, intensity values of H and M SILAC channels were divided by the summed up intensities of the Light channel (mock infected). Data was then averaged for label swap replicates and summed up for viral and host proteins independently. Finally, data was normalized to the 0-4h time period.

SILAC ratios of host proteins were processed by first transforming them into log2 space. The median SILAC H/L and SILAC M/L ratios from the input samples were used to estimate the mixing ratio of the input and the H/L and M/L ratios after the enrichment were adjusted correspondingly. SILAC H/M ratio that relate to the Pan/Mal (or Mal/Pan) infection treatment were normalized to 0. Then the replicate measurements were averaged. Proteins that were quantified in only one replicate were excluded. Then the top 2% of proteins (highest log2 fold-change) of either Pan/Mock or Mal/Mock condition were selected and multi-set GO-enrichment was performed using Metascape tool (http://metascape.org) ^74^.

Protein level data was matched to RNA level data based on the HGNC official gene symbol. Protein synthesis efficiencies were calculated by subtracting log10(RPKM) from log10(iBAQ) values.

For quantification of SILAC viral protein expression kinetics we extracted all quantifications of peptide level evidences for each individual replicate from evidence.txt. Median log2 ratios were then normalized to 0 for individual replicates. Comparative viral protein expression kinetics were based on Pan/Mal shared peptides. For each time period, replicate and viral protein the Pan/Mal SILAC protein ratio was calculated as the median of all SILAC peptide level ratios. Replicate SILAC protein rations were averaged and proteins only identified in one replicate were excluded.

For pSILAC, ratios for viral proteins other than M1 were extracted from proteinGroups.txt. M1 protein ratio was calculated based on shared peptides (see above). Non-normalized and log2 ratios were used in both cases. Quantifications of viral proteins with >75 % ratio variability were removed.

### RNA sequencing and data processing

Total RNAs from A549 cells with and without infection (infection conditions as described in the AHA-SILAC experiment) were extracted using TRIzol reagent (Life Technologies) following the manufacturer’s protocol. Truseq Stranded mRNA sequencing libraries were prepared with 500 ng total RNA according to the manufacturer’s protocol (Illumina). The libraries were sequenced on HiSeq 2000 platform (Illumina) and yielded in total 186 million 101-nt single-end reads. The sequencing reads were first subjected to adapter removal using flexbar with the following parameters: -x 6 -y 5 -u 2 -m 28 -ae RIGHT -at 2 -ao 1 -n 4 -j -z GZ ^75^. Reads mapped to the reference sequences of rRNA, tRNA, snRNA, snoRNA and miscRNAs (available from Ensembl and RepeatMasker annotation) using Bowtie2 (version 2.1.0)^76^ with default parameters (in --end-to-end & --sensitive mode) were excluded. The remaining reads were then mapped to the human and Pan/Mal influenza A reference genome using Tophat2 (v2.0.10) ^77^ with parameters: -N 3 --read-gap-length 2 --read-edit-dist 3 --min-anchor 6 --library-type fr-firststrand --segment-mismatches 2 --segment-length 26, and the guidance of RefSeq/Ensembl human gene structure and known viral gene annotation. Gene expression levels (RPKM) were estimated by Cufflinks (v2.2.1) ^78^ with parameters: -u --library-type fr-firststrand --overhang-tolerance 6 --max-bundle-frags 500000000. Splice junction reads for various M transcripts were counted with customized Perl scripts. Splice isoforms were accepted that had >500 read counts in both replicates.

### Transfections

A549 cells were seeded on 6-well plates and transfected with 2.5 μg of expression constructs and Lipofectamine 3000 (Thermo Fisher) reagent according to the manufacturer’s instructions. Cells were harvested by trypsinization and subjected to lysis and immunoblotting or qRT-PCR.

### Lysis, SDS Page and immunoblotting

Cells were lysed in lysis buffer (125 mM NaCl, 0.1% SDS, 1% NP-40, 5% glycerol, 50 mM Tris-HCl, pH 7.4) for 1 h on a rotating wheel and centrifuged. Supernatant was supplemented with NuPage LDS Sample buffer (Invitrogen), 50 mM DTT and heated for 10 min at 70°C. Samples were run on 4%–12% Bis-Tris gradient gels (NuPAGE, Invitrogen) before being blotted onto PVDF membrane (Immobilon-P, Millipore) using a wet blotting system (Invitrogen). Specific antibodies against the HA epitope (clone 3F10, Roche), vezatin (clone B-1, SantaCruz), M1 (clone GA2B, BioRad) or M2 (polyclonal, RRID: AB_2549706, Thermo Fisher) and suitable HRP-linked secondary antibodies were used.

### qRT-PCR

Cells were harvested by trypsinization and total cellular RNA was prepared, quality-controlled and reverse described. Prior to PCR, cDNA concentrations were adjusted to 2.5 ng/μl. Quantitative real-time PCR was performed on a 7500 Fast Real-Time PCR System (Life Technologies) using SYBR green as dye. The gene-specific primers for M1 were: fw 5′-CTAACCGAGGTCGAAACG-3′, rev 5′-CCCTTAGTCAGAGGTGAC-3′. For M2, forward primer were a 1:1 molar mixture of primers 5′-CTAACCGAGGTCGAAAC**T**CC-3’ and 5′-CTAACCGAGGTCGAAAC**C**CC-3′ and reverse primer: 5′-ACTCCTTCCGTAGAAGGCCC-3’. Ribosomal protein L32 (RPL32) was quantified with the forward primer: 5′-GATGCCCAACATTGGTTATGGA-3′ and the reverse primer 5′-GGCACAGTAAGATTTGTTGCAC-3′. The M1/M2 mRNA levels were normalized by subtraction of RPL32 threshold cycle (C_T_) values, resulting in ΔC_T_ values of the analyzed samples. ΔΔC_T_ values represent the fold-change between two samples in log2 space. 2^-ΔΔC_T_ values represent the values in non-log space. Data were analyzed using in-house generated R scripts and presented as means with standard deviations of triplicate samples.

### Immunofluorescence microscopy

A549 were grown on glass coverslips and infected with the indicated viruses at an MOI of 1 (FFU/cell). At the indicated time points postinfection, cells were fixed and processed for immunofluorescence staining essentially as described ^79^. Specific antibody against NP antigen was used (clone AA5H, Serotec). Nuclei were counterstained by 4′,6-diamidin-2-phenylindol (DAPI). Images were acquired by an Eclipse A1 laser-scanning microscope using the NIS-Elements software package (Nikon). At least 150 cells were counted per condition to quantify the subcellular distribution of NP using ImageJ software ^80^.

### Computational RNA structure analyses

M segment sequences were obtained from the NIAID Influenza Research Database (IRD) ^81^ through the web site at http://www.fludb.org. We used the following settings for human-adapted strains: date range >= 2009, sub-type H3N2, only complete genomes, include pH1N1 sequences, host human; exclude laboratory strains and duplicate sequences and geographic grouping: South America, Europe and Asia; for avian-adapted strains: date range >= 2009, only complete genomes, include pH1N1 sequences, host: avian; exclude duplicate sequences and geographic grouping: Europe and Asia. These two sets of sequences were merged with the respective references sequence, i.e. A/Panama/2007/1999 - Pan (human) and A/Mallard/439/2004 - Mal (mallard), and aligned using the program Muscle ^82^ resulting in two multiple sequence alignments (MSA), one for the human-adapted strains comprising 403 sequences and one MSA for the avian-adapted strains comprising 199 sequences. Both alignments are straightforward to establish based on the long open reading frame of the M1 isoform covering almost all of the M segment and on the overall high primary sequence conservation of the M segment. Evolutionary trees relating the sequences in either MSA were then derived using PhyML in conjunction with the HKY evolutionary model which provided the best fit to the data [version v3.0^83^].

These two input alignments (including the combined annotation of the known protein-coding M1 and M2 regions) and the corresponding evolutionary trees were then used as input to RNA-Decoder ^59^. We used RNA-Decoder to identify the RNA secondary structure that is best supported by the evolutionary signals contained in the two input alignments (the so-called maximum-likelihood structure). The predictions by RNA-Decoder also included the posterior base-pairing probabilities for each base-pair of the predicted RNA structure. Predicted base-pairs with a base-pairing probability smaller than 25% were omitted from the RNA structure visualization. Note that each multiple sequence alignment was analysed by RNA-Decoder in one chunk, i.e. without partitioning it artificially into sub-alignments.

Finally, the predicted RNA structure element nt 733-766 was plotted with the sequence of the Mal strain using VARNA tool ^84^ the RNA structures predicted for the two alignments of avian-and human-adapted sequences was visualized using R-chie ^85^ including information on the pairing probability of each base-pair.

### Nucleotide polymorphisms

For calculating percentages of nucleotide identities at the 3’ splice site we used the avian and human-adapted sequences as described above.

